# Reconstruction of critical-sized mandibular defects in a sheep model using a PLLA-PGA-CC scaffold

**DOI:** 10.64898/2026.06.25.734681

**Authors:** Katharina Pippich, Vincent Vitus Klett, Adem Aksu, Frank Reinauer, Stefan Milz, Andreas M. Fichter, Lucas M. Ritschl, Judith Reiser, Julia Werner, Christine Baumgartner, Achim von Bomhard

## Abstract

**Introduction:** Critical-sized bone defects cannot heal spontaneously, requiring additional, often burdensome, treatment. Thus, various synthetic substitute materials have been investigated regarding their treatment capacity. Poly-L-lactic acid (PLLA) and polyglycolic acid (PGA) have emerged as promising biodegradable scaffold materials. The addition of inorganic materials such as calcium carbonate (CC) has also been shown to be advantageous. This study investigates the effect on bone regeneration of PLLA-PGA-CC scaffolds in critical-sized bone defects over a two-year observation period using sheep as an animal model.

**Methods:** Critical-sized mandible angle defects were created in twelve female merino sheep. Mandibular defects were reconstructed with PLLA-PGA-CC scaffolds in four sheep, while the remaining eight served as negative control (defects left empty). The scaffolds were manufactured using computer-aided design and manufacturing, incorporating an interconnected porous structure and fixated with polyether ether ketone cages. Bone regeneration was evaluated using computed tomography (CT) imaging at 3, 12, and 24 months postoperatively. Bone volume was assessed quantitatively. Additionally, a histological analysis was performed.

**Results:** Surgical procedures were successful and without major complications. CT assessment showed more bone regeneration in the scaffold group (mean volume: 7,472 mm³) than in the control group (4,168 mm³, *p* = 0.1) at 24 months postoperatively. Resorption of the scaffolds and formation of compact lamellar bone tissue were confirmed by histological analysis. However, the osteoconductive properties of the scaffolds were limited, with only minimal ingrowth of bone tissue into the porous structure. In both groups, fibrous tissue infiltration and the formation of cyst-like cavities in the defect region were observed.

**Conclusion:** PLLA-PGA-CC scaffolds were found to be biocompatible and enhanced bone regeneration compared to the control group. Due to fibrous tissue infiltration and the lack of osteoconductivity, the suitability of the material for critical-sized bone defect reconstruction is limited.

## Introduction

Bone tissue is known for its remarkable regenerative potential, but large defects do not heal spontaneously [1]. Critical-sized bone defects are generally defined as those that don’t heal spontaneously or by means of surgical fixation and are associated with significant burden and clinical challenges [1, 2]. Thus, critical bone defects usually require some kind of bone substitution. They are often the result of infections, osteomyelitis, tumor resections, trauma injuries, or non-union fractures [2, 3]. Defects can be bridged using substitute materials or bone grafting. Tissue-engineered scaffolds are in clinical use today, but autologous transplantation is still the gold standard for critical-sized defect reconstruction [4, 5]. Although autografts generally have good biological properties and do not usually elicit an immune response, they have several drawbacks. These include additional defects at the bone harvesting site, prolonged duration of the surgical procedure, and increased morbidity [6]. Furthermore, the amount of bone is limited in quantity and possible shapes, and the graft must be manually adapted to the defect. Current research therefore focuses on synthetic scaffolds and bioactive substances, such as growth factors and cell seeding [7, 8]. These scaffolds should be biocompatible, osteoconductive, bioactive, and osteogenic to improve regenerative capabilities. To promote bone regeneration, as well as act as a carrier for cells and bioactive agents, scaffolds with an interconnected porous structure are the preferred option [9, 10].

Biodegradable polymers have emerged as a promising material for bone tissue engineering [11]. Poly-L-lactic acid (PLLA) and polyglycolic acid (PGA) are often used as scaffold materials, possessing adequate mechanical properties without producing harmful by-products [12, 13]. Often, different polymers are combined to achieve ideal properties, such as the rate of degradation [14]. Furthermore, the addition of inorganic materials like calcium carbonate (CC) is known to be advantageous. Calcium carbonate balances acidic milieus caused by scaffold degradation products and promotes bone formation [15].

In this study we investigated porous 3D-printed PLLA-PGA-CC scaffolds for the regeneration of critical-sized sheep mandible defects. In previous in vitro testing this material showed superior results compared to a variety of polymer-based materials [16]. As most current studies are limited by the utilization of less-suited small-animal models and short observation periods, we used a sheep model over a two-year period [17]. A cell-free approach was chosen, as cell harvesting and seeding are associated with regulatory concerns, additional effort, high cost, and further risks, and inhibit the transition from the bench to the bedside [18, 19].

## Materials and methods

### Animals

Twelve adult merino sheep were procured from a local breeder (Schafzucht Schleich, Brunnthal, Germany). Sheep were chosen because of their anatomical similarity to humans [3]. The sheep were uniformly sized, female, and aged around six months, with a mean weight of 63.3 kg (51–78.8 kg). Animals were acclimatized at least three days before surgery. In total twelve sheep (unilateral mandibular angles) were included in the study. These were divided into two groups. The study group (n=4) was implanted with PLLA-PGA-CC scaffolds. In the remaining sheep (n=8), mandibular defects were not bridged by scaffolding in order to serve as a negative control.

The study was carried out in accordance with European guidelines for the care and use of laboratory animals and ARRIVE guidelines [20]. The animal experiment was approved by the local administrative body (Regierung von Oberbayern, registration number: ROB 55.2-2532.vet_02-18-75).

### Manufacturing of the scaffolds

Multiple bioresorbable polymers were tested in vitro beforehand. Because of the promising results, poly-L-lactic acid–polyglycolic acid–calcium carbonate (PLLA-PGA-CC) was chosen for in vivo testing [16]. Scaffolds were manufactured using computer-aided design and manufacturing (CAD/CAM) and packaged and gamma-sterilized by KLS Martin SE & Co. KG (Tuttlingen, Germany) under industrial clean-room conditions. Polyether ether ketone (PEEK) cages for scaffold fixation, placeholders for later cell seeding possibilities, cutting guides, were also manufactured by KLS Martin and sterilized prior to surgery. The PEEK cages were manufactured for mandibular fixation of the scaffolds with osteosynthesis screws (MaxDrive, KLS Martin). The PEEK cage distributes the force exerted on the scaffold over a wider surface. Precise implantation and consistent creation of defects were ensured by using guides for cutting the mandibular angles and pre-drilling for fixation screws. The component design was based on computed tomography (CT) scans of one sheep, using CAD. 3D-printing of the scaffolds allowed for a porous structure with interconnected channels (Figure 1). Pores (800 μm) and interconnectivity were intended to achieve a trabecular structure similar to physiological bone, as well as guiding the ingrowth of bone tissue. Furthermore, the microporous structure of the material aimed to facilitate cell adhesion. The scaffolds were additively manufactured using FDM technology. For 3D-printing an ARBURG AKF freeformer 200-3X (ARBURG, Germany) was used.

**Figure 1:**
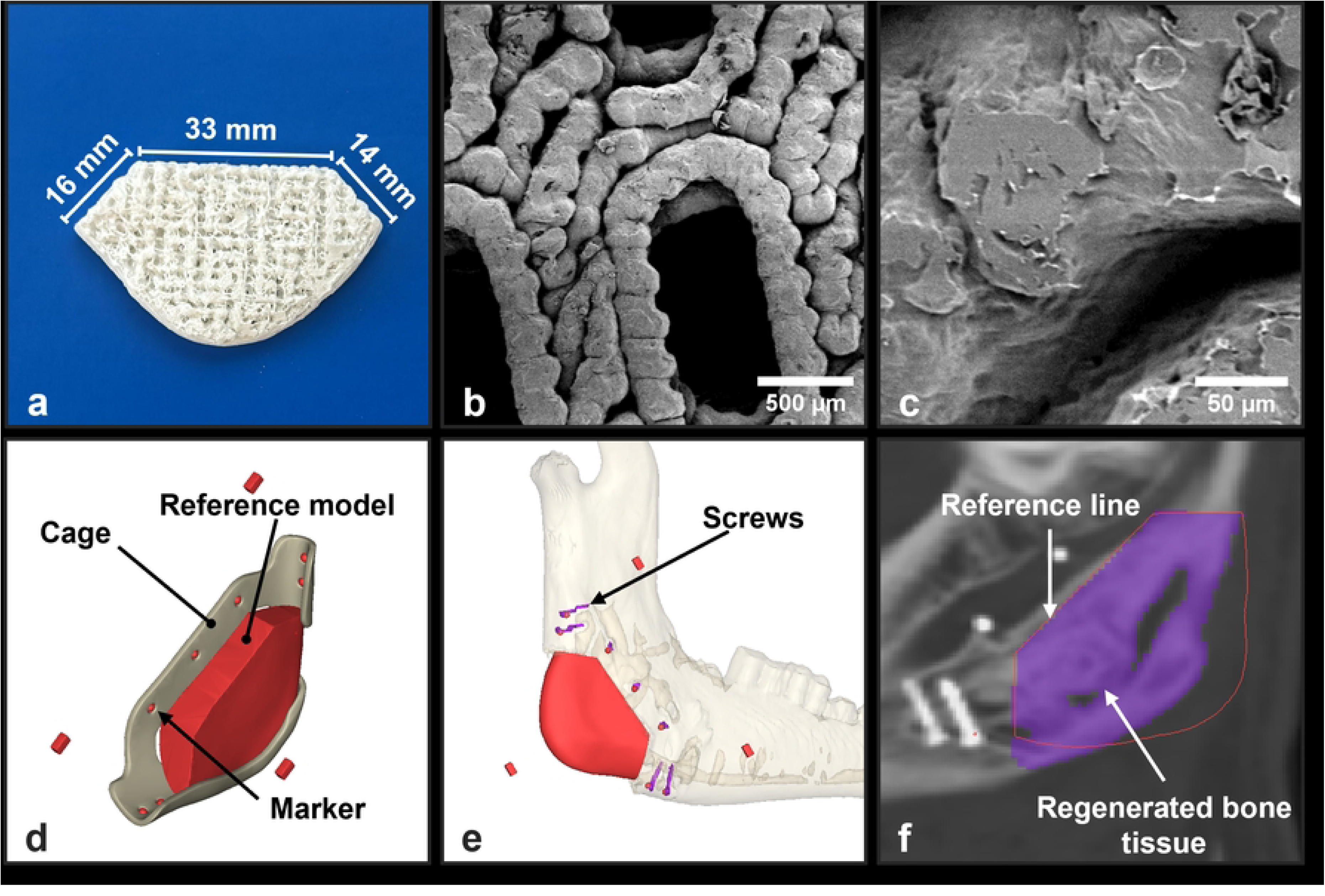
**Scaffold structure (a–c): (a)** 3D-printed scaffold with dimensions; **(b)** REM image of pores (50× magnification); **(c)** REM image of micro-porous structure (500 x magnification); **Quantitative CT assessment (d–f): (d)** Reference model with markers relative to screw holes in the PEEK cage; **(e)** STL model with screw location markers positioned by alignment with segmented screws; **(f)** Region of interest established by outline of the scaffold STL model; Segmentation of regenerated bone tissue (purple).

### Anesthesia

After intramuscular application of xylazine (approx. 0.1–0.5 mg/kg) and subsequent intubation, anesthesia was induced by bolus injection of propofol 2% (approx. 4–8 mg/kg) under mechanical ventilation. Metamizole (approx. 25–50 mg/kg) and ketoprofen (approx. 3 mg/kg) were administered simultaneously. Anesthesia was maintained with isoflurane (approx. 1.8–2%) under intermittent positive pressure ventilation. Intraoperatively, ketamine (approx. 8 mg/kg/h) and fentanyl (approx. 0.001–0.01 mg/kg, every 20–30 minutes) were administered intravenously.

### Surgical procedure

All surgical procedures were carried out under sterile conditions. Sheep were shaved and washed under general anesthesia. The surgical field was disinfected. In a supine position with the head extended, a submandibular incision (approx. 15 cm) was made medially along the mandible body. Then the external jugular vein was exposed and lateral branches obstructing access to the mandibular angle coagulated or ligated and separated. The jugular vein was then pushed in a lateral direction. Periosteum and muscle insertions were detached from the inner and outer sides of the mandible with a raspatory. Then the caudal section of the mandible, including the mandibular angle, was exposed and a bone defect (30 x 30 mm, trapezium-shaped) was created. The defect was cut along the 3D-printed cutting guide with an oscillating saw caudally/distally from the mandibular canal. The scaffold was then placed in the bone defect. A PEEK cage covering the mandibular angle was fixated with osteosynthesis screws. This technique allows for fixation without the screws being placed in the scaffold itself. In addition, the screws served as markers, enabling reliable identification of the defect area in 3D imaging (Fig 1F). Wound closure followed with resorbable suture subcutaneously (3–0) and intracutaneously (4–0).

In the control group, mandibular defects were created, as described above, and the PEEK cage was fixated without scaffolds. Through the implantation of the PEEK cage at the empty mandibular defect, comparable conditions were assured.

### Postoperative management

Buprenorphine (0.005–0.02 mg/kg) was administered for at least two days after reconstruction surgery every 8–12 hours. Administration was continued if signs of pain were identified. After mandibular reconstruction a regeneration period of at least 40 days was ensured before the first CT scan.

### Computed tomography assessment

CT scans were conducted under general anesthesia at 3, 12, and 24 months postoperatively (0.9 mm slice thickness; 0.45 mm slice increment). Scans were stored in DICOM format. For the analyses, Materialise Mimics Medical Software (Version 23.0.2.530, Materialise NV ©) was used. Qualitative analysis was performed to assess bone regeneration and morphology macroscopically.

The volume of newly formed bone in the study group and control group over the two-year timespan was determined quantitatively. In the first step, solid 3D STL models of the scaffolds were imported into the CT files in Mimics for the bilateral mandibular angles. These models have markers at the screw positions on the PEEK cage (Figure 1F). Then the screws in the CT scans were segmented and the scaffold models were positioned by aligning the markers and screws. This allowed for a reference to the originally cut-out defect (i.e., region of interest) in the follow-up scans, to distinguish newly formed and native bone and for ensuring systematic evaluation. The positioning of the scaffold models was precise at the cut line of the defect, but precise positioning laterally and medially was not possible. Newly formed bone within the scaffolds (empty defect in the control group) and lateral/medial to the scaffold was therefore included. Bone volume was determined by reviewing every slice within the region of interest and marking newly formed bone. The marker was always set to a threshold of 226–3,071 Hounsfield units (HU), to prevent the marking of non-bone tissue. Then 3D models of the segmentations were calculated. As PLLA-PGA-CC is radiolucent, scaffold volume could not be quantified [21].

### Histological assessment

Mandibular angle specimens were fixed and then dehydrated in ascending fractions of alcohol (70%, 80%, 90%, 100%). Fat was dissolved using xylene, and specimens were embedded in methyl methacrylate (MMA, Fluka, Switzerland). Controlled polymerization was carried out after the methyl methacrylate fully penetrated the specimens, using benzoyl peroxide as a starter. Sections were cut (sections along the mandibular angles’ centers) using a diamond coated band equipped band saw (cut-grinder, Walter Messner GmbH, Germany). At this point, contact radiographs (Faxitron, Faxitron X-ray Corporation, USA) with AGFA Structurix X-ray sensitive films (AGFA, Germany) of the specimens’ sections were obtained. Prior to staining, the sections were glued onto plastic slides, ground, and polished (400CS, EXAKT Advanced Technologies GmbH, Germany). The surfaces of the thin sections were then etched in formic acid for 15 minutes and stained with Giemsa solution (Sigma Aldrich) for 30 minutes at 60°C. After dipping in acetic acid and rinsing, eosin (0.1%) and glacial acetic acid (20 µl) were applied for 4 minutes.

Stained thin sections were analyzed using an Axiophot (Zeiss, Germany) transmitted-light microscope with Plan-Neofluar (Zeiss, Germany) objectives (5 ×, 10 ×, 20 ×). Magnified images were taken with an Axiocam HRc (Zeiss, Germany) digital camera in conjunction with Zeiss Axiovision® imaging software. Additionally, stitched images of the sections were created using an Olympus BX51 microscope (Olympus, Japan) attached to a 2D scanning table (Ludl Electronic Products Ltd, USA) and 4 × objective. For stitching, Stereo Investigator (MBF Bioscience, USA) software was used. Stitched files were viewed using the Biolucida viewer (MBF Bioscience, USA).

### Statistical analysis

The results of each group are presented in mean values ± standard deviation. Statistical analyses were performed in Python (version 3.10.17) with the packages pandas, numpy, scipy.stats, and scikit-posthocs. To analyze differences between groups at a single time point, the Kruskal–Wallis test was applied. When significant, pairwise post-hoc comparisons were conducted using Dunn’s test with Bonferroni correction. The significance level was set at α = 0.05.

## Results

Surgical procedures were successful and there were no intraoperative or immediate complications. All animals were returned to feeding the same day as the surgery. The sheep showed no signs of pain, and no weight loss occurred. Soft tissue thinning caused by atrophy resulted in non-inflammatory exposure of one mandibular angle. Due to animal welfare concerns the affected sheep was euthanized 12 months after surgery. The 24-month results apply to eight mandibular angles, three of the scaffold group and five of the control group.

### Computed tomography assessment

Bone regeneration occurred in all mandibular angles (Fig 2). Overall, more pronounced bone formation was observed in the polymer scaffold group, particularly at the final follow-up, compared to the control group. At 3 months, CT images showed only limited ingrowth of bone tissue into the porous scaffold structure. Bone formation was mainly located lateral to the scaffold, with new bone extending over the PEEK cage toward the resected area, especially at the edge of the mandibular angle.

**Figure 2:**
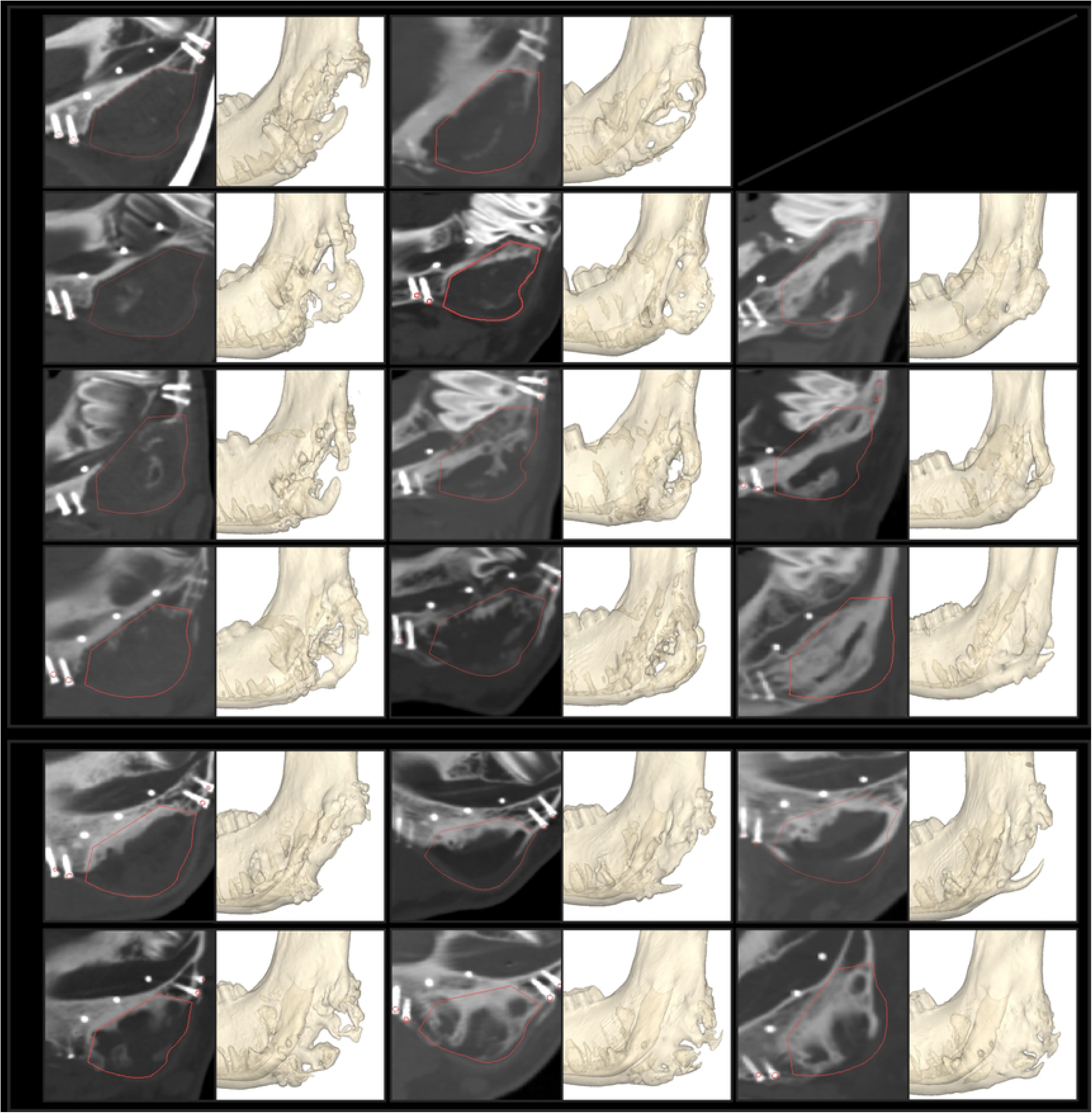
2D and 3D images of computed tomography assessment at 3, 12, and 24 months postoperatively. Scaffold group (S1–S4); Control group representative images (S7, S11). Red lines for initial defect reference.

At 12 months, increased bone formation originating from the surgical margins was visible within the scaffold region. Nevertheless, most new bone tissue continued to form laterally and medially to the scaffold. The bone appeared to grow in a branch-like pattern. After 24 months, further bone tissue had developed within the scaffold area. Branches of newly formed bone arising from the mandibular ramus and body converged at the angle and partially merged with bone extending from the surgical margin. Although the defects showed substantial regeneration, residual gaps—particularly in the central region—resulted in heterogeneous bone morphology.

In the control group, defects did not fully regenerate and gaps or cavities remained visible (Fig 2). Bone predominantly grew from the surgical margin, becoming thinner toward the angle. Additionally, less directed bone formation occurred medially and laterally, as well as along the PEEK cage. Overall, bone regeneration patterns in the control group were heterogeneous. In some cases, the defect area was largely bridged, but the resulting bone was thin and frequently exhibited discontinuities.

Quantitative analysis demonstrated increasing bone volumes within the resected area over time in all sheep (Table 1). At 3 months, mean bone volume measured 1,842 mm³ (± 785 mm³ SD) in the scaffold group and 2,131 mm³ (± 779 mm³ SD) in the control group. By 12 months, bone volume increased to 4,859 mm³ (± 2,310 mm³ SD) in the scaffold group and 3,330 mm³ (± 1,249 mm³ SD) in the control group. At 24 months, mean bone volume reached 7,472 mm³ (± 244 mm³ SD) in the scaffold group and 4,168 mm³ (± 2,070 mm³ SD) in the control group. Differences between groups were not statistically significant at any of the follow-ups (*p* = 0.85; 0.27; 0.10, respectively).

**Table 1:** Results of the quantitative computed tomography assessment: Volume of bone formation in the resection region; Sheep 5 excluded due to complications.

| Follow-up: | Bone volume mm<sup>3</sup> |  |  |
| --- | --- | --- | --- |
|  | 3 months | 12 months | 24 months |
| **Scaffold group** |  |  |  |
| Sheep 1 | 970 | 1,951 | - |
| Sheep 2 | 1,918 | 4,096 | 7,231 |
| Sheep 3 | 1,621 | 7,040 | 7,465 |
| Sheep 4 | 2,860 | 6,348 | 7,719 |
| Mean (± SD) | 1,842 (± 785) | 4,859 (± 2,310) | 7,472 (± 244) |
| **Control group** |  |  |  |
| (Sheep 5) | - | - | - |
| Sheep 6 | 3,626 | 5,986 | 7,576 |
| Sheep 7 | 1,511 | 2,477 | 3,227 |
| Sheep 8 | 1,721 | 2,987 | 3,310 |
| Sheep 9 | 1,585 | 3,206 | - |
| Sheep 10 | 1,817 | 2,142 | 2,220 |
| Sheep 11 | 1,891 | 3,382 | 4,505 |
| Sheep 12 | 2,766 | 3,130 | - |
| Mean (± SD) | 2,131 (± 779) | 3,330 (± 1,249) | 4,168 (± 2,070) |

### Histological assessment

Thin sections showed full degradation of the scaffold. In sheep 2 and 4 the resected zone was mostly filled with compact bone (**Figure 3**). Bone was lamellar in structure. The bone tissue showed signs of remodeling. Osteons and pockets of bone marrow traversed the newly formed bone tissue. In the thin section of sheep 3, a cyst-like cavity (11 x 19 mm), presumably containing fluid and connective tissue, was found. In the surrounding area, rudimentary bone formation consisting of cancellous and compact bone occurred. Further connective tissue throughout the region of interest was visible. Sheep 2 also had multiple cyst-like cavities (1 x 2 mm, 4 x 6 mm, 6 x 13 mm) within the regenerated bone. In sheep 4 there were two cavities filled with connective tissue in the center of the defect. Radiographs of the thin sections and resected specimen depicted mostly compact bone, with regions of cancellous bone tissue. Gaps were visible where connective tissue ingrowth appeared.

**Figure 3:**
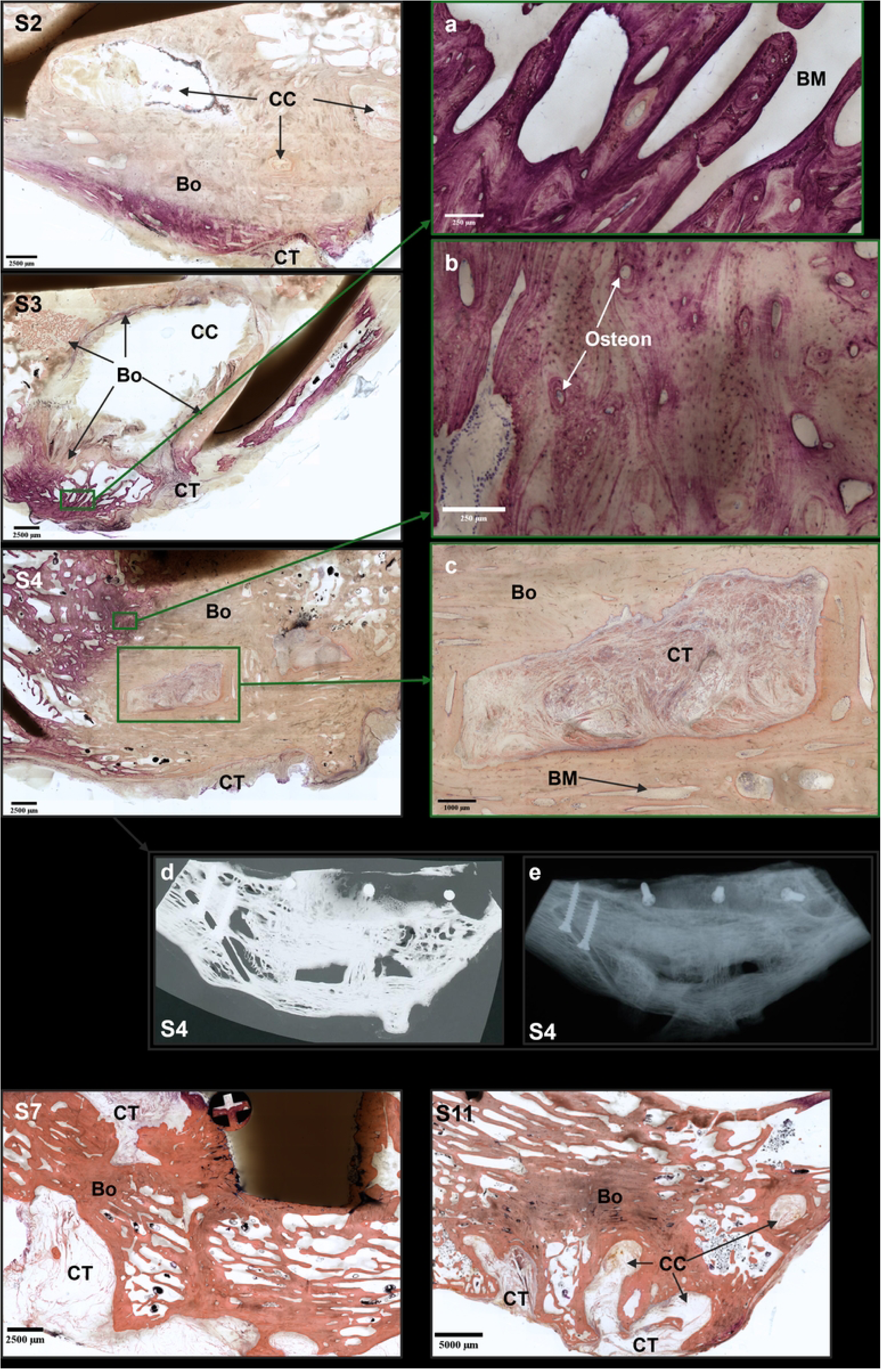
Histological assessment of thin sections: **Sheep 2 (S2):** Compact bone tissue in resected area (Bo) bordered by connective tissue (CT) at the angle. Non-inflammatory cyst-like cavities (CC) filled with fluid and connective tissue; **Sheep 3 (S3):** Large non-inflammatory cyst-like cavity in defect zone surrounded by compact and trabecular bone tissue. Connective tissue within bone and at the angle; **Sheep 4 (S4):** Mostly compact bone tissue in the defect area, bordered with connective tissue along the angle. Centrally two connective tissue cavities are embedded in the bone tissue; **Sheep 7 (S7):** Mostly lamellar trabecular bone tissue within the defect. Bone tissue is bordered by connective tissue; **Sheep 11 (S11):** Lamellar trabecular, centrally compact, bone tissue in the resected zone. Several non-inflammatory cystic cavities within bone tissue; **(a):** Trabecular bone tissue with remodeled lamellar structure. Bone marrow (BM) in between trabeculae; **(b):** Lamellar structure of compact bone tissue. Signs of remodeling, osteons are visible throughout the bone tissue; **(c):** Connective-tissue-filled cavity in compact bone. Pocket of bone marrow in the compact bone; **(d):** Contact radiography of thin section (S4): Mostly compact bone with areas of trabecular bone. Radiolucency in the center; **(e):** Contact radiography of resected mandibular angle specimen (S4): Mandibular angle vastly regenerated with gaps centrally; bone (Bo), cyst-like cavity (CC), connective tissue (CT), bone marrow (BM).

Cystic cavities were also present in specimens of the control group (**Figure 3**). These cavities were filled with fluid or connective tissue. In one case, muscle tissue was identified within the bone tissue. In this specimen, osteomyelitis had occurred. In the control group, defects were not fully regenerated. The cross sections showed bone formation not reaching the outer edge of the mandible angle. The bone tissue was bordered by connective tissue. In these specimens, connective tissue was also found within the defect area.

## Discussion

In this study, we evaluated a novel polymer scaffold material for the reconstruction of critical-sized mandibular defects in a long-term sheep model. Additive CAD/CAM-based manufacturing enabled the production of individualized scaffolds, PEEK cages, and surgical guides. Precise preoperative planning together with cutting and drilling guides facilitated accurate scaffold placement and reduced operative time, as previously demonstrated [22]. No major complications occurred in the scaffold group, and neither implant encapsulation nor rejection was observed, underscoring the material’s biocompatibility.

Overall, bone volume within the resected areas was greater in the scaffold group compared to controls (Fig 3). At 3 months, bone volumes were slightly lower in the scaffold group, although this difference was not statistically significant (14 ± 42% SE, *p* = 0.85) (Fig 4). At 12 months, the scaffold group showed more newly formed bone tissue, again without significant difference (46 ± 27% SE, *p* = 0.26). After 24 months, mean bone volume further increased in the scaffold group; however, the difference remained statistically non-significant (79 ± 25% SE, *p* = 0.10). Although the differences did not reach significance at any time point, the p-values decreased over the study duration. This may be explained by high variance and small sample size. Due to non-normal data distribution, non-parametric statistical methods were applied. Regarding the quantitative CT analysis, minor variability in the positioning of the reference model when establishing a consistent region of interest is to be expected, though we assume this did not meaningfully influence results. Threshold-based segmentation minimized the likelihood of human error in identifying osseous tissue.

**Figure 4:**
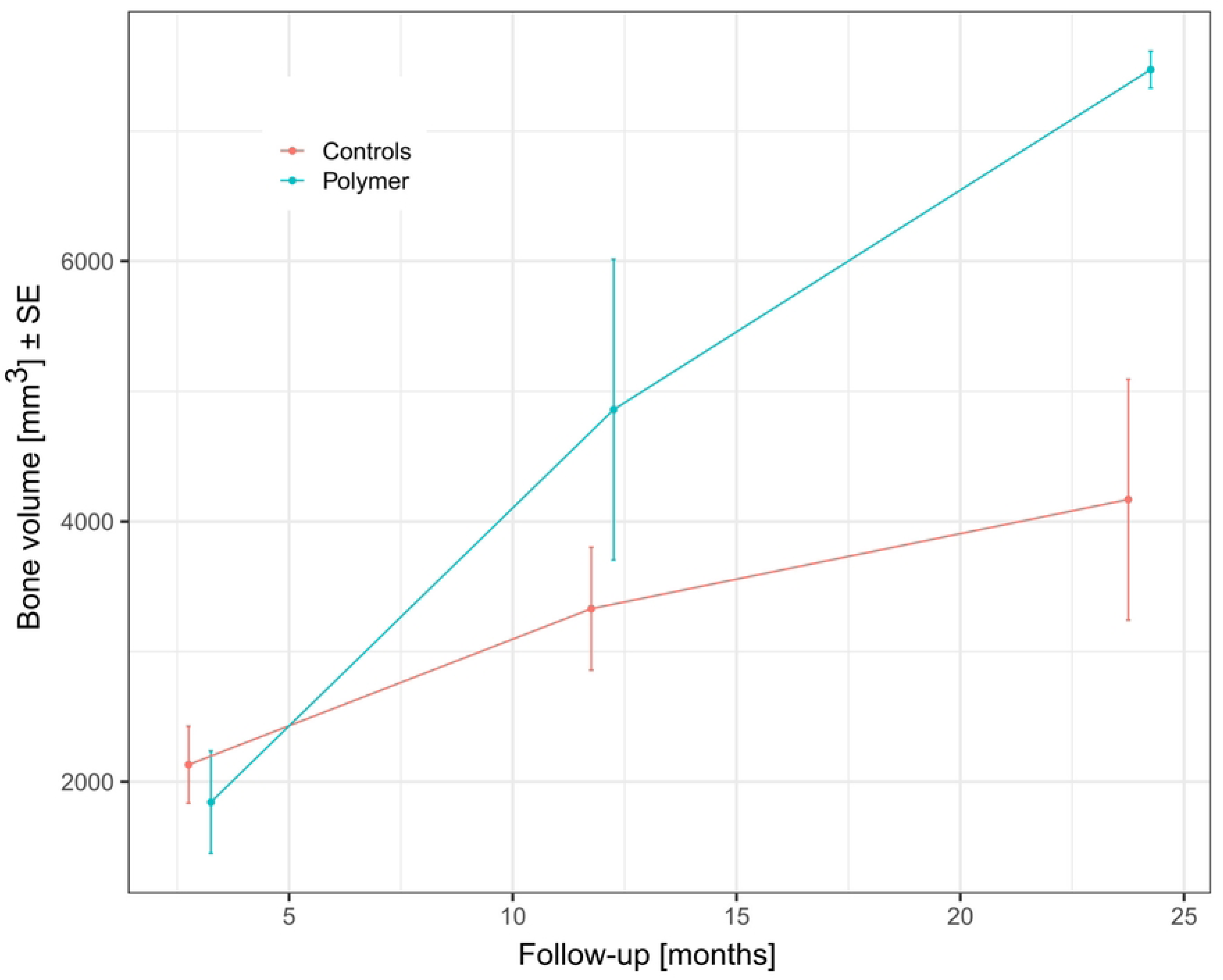
Comparison of bone volume (± standard error) in the scaffold group (polymer) and the negative control at 3, 12, and 24 months postoperatively.

Despite lesser bone regeneration in the control group, the overall extent of bone formation remained unexpectedly high. We attribute this to the presence of the PEEK cage combined with periosteal coverage, which likely biased outcomes. The PEEK cage functioned as a physical barrier preventing soft tissue collapse into the defect and provided structural stabilization that may have promoted bone formation even without a scaffold. Additionally, the periosteum is well known for its osteogenic capacity, and bone healing failure in critical defects is often associated with its absence [23, 24]. Thus, periosteal preservation in combination with cage stabilization may have guided bone formation. This is consistent with the observed pattern of initial bone apposition predominantly lateral and medial to the defect rather than centrally, suggesting periosteal-driven appositional growth. Bone also appeared to form along the PEEK cage, particularly at the rounded posterior portion of the cage at the mandibular angle (Fig 2, S7), where a branch-like bone pattern was evident.

Histological evaluation demonstrated good microscopic bone quality in both groups, with lamellar structure and ongoing remodeling. More compact bone formation was evident in the scaffold group (with the exception of sheep 3), whereas the control group exhibited a predominance of cancellous bone (Fig 3). Macroscopically, regenerated bone in both groups remained inadequate. Connective tissue was present within and at the margins of the regenerated bone, indicating incomplete osteogenesis within the scaffold and early fibrous infiltration prior to bone deposition. In controls, connective tissue surrounded newly formed bone, limiting further growth and preventing complete defect regeneration—an expected finding in critical-sized defects where fibrous tissue occupies the empty space. This may explain the superiority of bone formation in the scaffold group: although the material lacked osteoconductivity, the scaffold served as a placeholder, maintaining space for ossification during degradation [25]. Cyst-like cavities were visible in CT scans and histological thin sections in both groups (Fig 2, Fig 3).

The PLLA-PGA-CC scaffold demonstrated insufficient osteoconductivity. At 3 months, virtually no bone ingrowth was detected within its porous structure. Other studies similarly report limited bone formation in PLLA-based implants [26]. Bone formed mainly along the scaffold periphery. By 12 and 24 months, bone originating from the resection margin extended into the former scaffold region. This suggests that bone formed only after scaffold degradation rather than within the material’s grid structure before resorption. PLLA-based materials are known to generate acidic degradation products that lower local pH [11, 27]. Although calcium carbonate was incorporated to buffer acidity, it may not have sufficiently stabilized the environment [15, 25, 28, 29]. This potentially contributed to poor osteoconductivity [30].

Due to the radiolucency of polymeric scaffolds, in-vivo degradation could not be assessed radiographically. However, histological analysis after 24 months confirmed complete scaffold resorption.

## Conclusions

In this study we tested a PLLA-PGA-CC scaffold in critical-sized bone defects in a two-year sheep model. The material is biocompatible and there was no encapsulation or rejection of the scaffolds. There was more bone formation in the scaffolded defects compared to non-bridged defects of the negative control group (*p* = 0.1). Both in the scaffold and control groups, cavities and connective tissue formed in the defect area. Furthermore, the osteoconductive properties of the polymer scaffold were insufficient. We therefore conclude that PLLA-PGA-CC is not ideal as a scaffold material for critical-sized bone defects.

## Acknowledgements

We gratefully acknowledge the support provided by Claudia Wittner and Thomas Potrusil (CADS GmbH, Austria), who performed the statistical analysis.

We used AI-based assistance (Microsoft Copilot) solely for linguistic support, including improving wording, clarity, and internal consistency during manuscript writing. All scientific content—such as data analysis, interpretation, and the formulation of conclusions—was conceived, evaluated, and confirmed entirely by the authors. No generative AI system was employed to produce, process, or interpret any scientific data.

